# Structural, Thermodynamic, and Dynamic Descriptors for Differential Mechanism of HIF-2 Activity Modulators

**DOI:** 10.1101/2025.04.23.650354

**Authors:** Rajat Punia, Gaurav Goel

## Abstract

Hypoxia-inducible factors (HIFs) are heterodimeric transcription factors critical for cellular adaptation to low oxygen conditions. Both enhancement and inhibition of HIF-2 activity has been shown to have a significant therapeutic relevance. We used all atom molecular dynamics (MD) simulations to elucidate the molecular mechanism for the differential effects of HIF-2 ligands on its complex with aryl hydrocarbon receptor nuclear translocator (ARNT). We find that agonists and antagonists differentially alter the binding site conformation, enthalpically strengthening or weakening the heterodimerization, respectively. These local changes were linked to conformational fluctuations of the HIF-2/ARNT complex, adding an entropic component to the effect of ligand binding on stability. Finally, we report a ligand-dependent effect on the DNA-binding bHLH domain’s structure and dynamics, impacting the hypoxia response element (HRE) binding. Based on these findings, we present a multiscale approach to enable high-throughput virtual screening of ligands for differential modulation of the structure, dynamics, and transcriptional activity of the HIF-2/ARNT complex.

**TOC Graphic:** 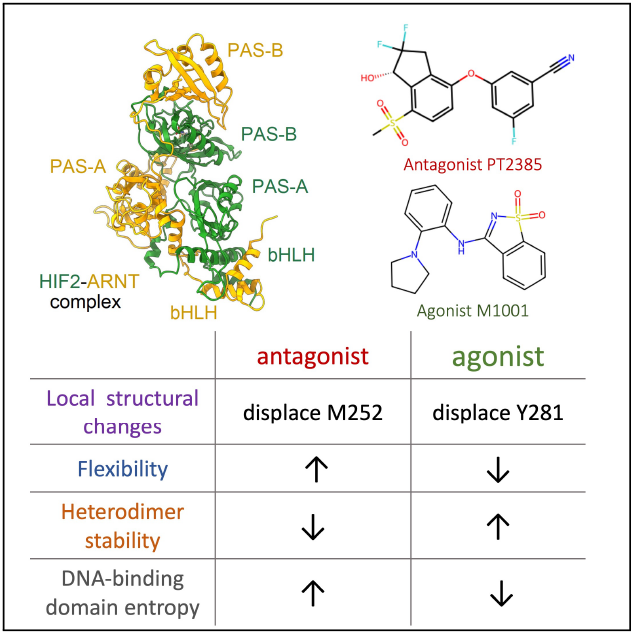

## Introduction

Hypoxia-inducible factors (HIFs), which include HIF-1*α*, HIF-2*α*, and HIF-3*α*, are a group of heterodimeric transcription factors that bind to hypoxia response elements in the genome, initiating signaling cascades that help cells adapt to low oxygen conditions (hypoxia). In normal oxygen conditions (normoxia), HIFs are rapidly degraded by prolyl hydroxylase domain (PHD) enzymes through oxygen-dependent proteasomal degradation. ^1–3^ However, the degradation of HIFs is inhibited under hypoxic conditions, allowing them to form heterodimers with aryl hydrocarbon receptor nuclear translocator (ARNT or HIF-1*β*), which in turn activate specific gene expression programs.^4,5^ These gene programs enhance oxygen supply and nutrient delivery to cells by promoting angiogenesis and erythropoiesis, which prevent cell death.^6,7^ HIFs also increase the expression of glucose transporters and glycolytic enzymes, which support the metabolic needs of cells in hypoxic environments.^8^

Hypoxia is a common feature in solid tumors due to high oxygen demand of rapidly proliferating cells.^9^ Tumors experiencing increased levels of hypoxia tend to be more resistant to therapy, have a higher probability of metastasis, and are associated with a poorer prognosis. ^7,10^ This leads to a strong association between increased HIF activity and cancer pathogenesis and progression.^11^ Tumors driven by deficiency of the Von Hippel-Lindau tumor suppressor (VHL) gene product, which is involved in HIF-2*α* degradation, are natural candidates for targeted inhibition of the hypoxia pathway.^12^ Stabilized HIF-2*α* promotes the expression of target genes involved in cell proliferation, survival, and angiogenesis, contributing to tumor growth and progression. ^10^ Belzutifan,^13^ a first-in-class HIF-2*α* specific inhibitor, recently received FDA approval for the treatment of nonmetastatic renal cell carcinomas, pancreatic neuroendocrine tumors, and central nervous system hemangioblastomas in patients with von Hippel–Lindau (VHL) disease.^14^

Specific activation of HIF-2 has also been investigated for its therapeutic potential, particularly in the treatment of renal anemia associated with chronic kidney disease (CKD), where it plays a crucial role in regulating erythropoietin (EPO) production. ^15–18^ Specific activation of HIF-2 is also desirable in conditions such as acute kidney injury (AKI),^19^ which is a critical factor in the onset of multiple-organ failure.^20^ Current therapeutics, such as HIF-prolyl hydroxylase (HIF-PH) inhibitors, ^21–25^ upregulate HIF activity but lack selectivity, leading to increased levels of all three HIF subtypes and causing side effects.^26^ For example, in 2021, the US FDA advisory committee recommended against approving Roxadustat due to a strong thrombosis signal observed in its clinical trials. Direct HIF-2*α* agonists could provide a more targeted approach, specifically enhancing EPO expression and potentially reducing side effects. When used in combination with HIF-PH inhibitors, these agonists could improve both the activation potential and specificity for HIF-2*α*,^27,28^ potentially leading to more effective and safer treatments.

The discovery of drugs for HIF activity modulation has remained challenging due to the complex nature of HIF signaling pathways and the difficulty in achieving specificity for individual HIF subtypes. Pioneering studies by Scheuermann et al. have focused on the HIF-2 PAS-B domain cavity, exploring its potential for allosteric modulation of HIF2-ARNT stability and HIF-2 specific transcriptional activity.^29–31^ These studies have led to significant breakthroughs, including the discovery of potent HIF-2 antagonists such as PT2977 (Belzutifan).^13^ Additionally, weak HIF agonists, particularly benzoisothiazole derivatives, have been identified, demonstrating synergistic effects when used in combination with approved HIF-PH inhibitors for treating renal anemia. ^27,28^

In this study, we used all atom molecular dynamics (MD) simulations to elucidate the underlying mechanism for the differential effects of agonist and antagonist binding on HIF-2 activity. Specifically, we have worked on identifying computational descriptors to advance the mechanistic understanding of effect of ligand binding at the PAS B domain of HIF-2 in the heterodimeric complex. Further, we present a multiscale approach based on these descriptors to allow separation of agonists, antagonists, and decoys, and thus, enable high-throughput in silico ligand screening for modulating structure and dynamics, and as a result, transcriptional activity of the HIF-2/ARNT complex. Toward this end, we present a set of MD simulations derived descriptors, including: (i) local conformational changes at the ligand-binding site, (ii) global fluctuations and binding affinity of the HIF2-ARNT complex, and (iii) allosteric changes in the structure and dynamics of the DNA-binding domain.

The HIF-2 transcription factor functions as a heterodimer consisting of HIF-2*α* and the constitutively expressed ARNT. This dimerization results in an asymmetric quaternary structure, with the basic helix-loop-helix (bHLH) domains of HIF-2*α* and ARNT forming a DNA-reading head that specifically binds to hypoxia response elements (HREs) in target genes (Figure 1A). The discovery of HIF-2 activity modulators has historically been challenging until the identification of an unusually large buried cavity within the HIF2*α* PAS-B domain, which is occupied by eight ordered water molecules in the apo-state.^29^ This cavity is analogous to the cofactor binding sites found in PAS domains of signaling proteins, such as the aryl hydrocarbon receptor (AHR), where cofactor binding allosterically regulates transcriptional activity.^32^ To investigate the plasticity of this cavity, we performed three independent 500 ns (total 1.5 µs) long MD simulations of HIF2-ARNT complex in ligand-unbound (apo) state using GROMACS 2021.4^33^ with the CHARMM36 force field.^34^ The initial structure of complex was taken from Protein Data Bank (PDB id 4ZP4^35^). The system was solvated in TIP3P water with 0.1 M NaCl, and equilibrated under NVT and NPT conditions at 310 K and 1 bar using the velocity-rescale thermostat and C-rescale barostat (complete details in Supporting Information Sec. 1.2). Figure 1B shows the distribution of HIF2*α* PAS-B cavity volume, calculated using POVME 3.0,^36^ highlighting the inherent flexibility of the cavity. The cavity volume ranged from 50 Å^3^ to 200 Å^3^, and these substantial variations are primarily due to side-chain reorientation of the cavity residues, as the C_*α*_ RMSD of these residues relative to the initial experimental apo state remained very small (approximately 1 Å) throughout the simulation. Targeting this flexible PAS-B domain cavity has led to the discovery of several HIF-2 antagonists^13,27,29,30,35,37^ and weak agonists.^27,28^ As the PAS-B cavity adopts a range of volumes across known ligand-bound (holo) structures (see Figure 1B), incorporating this flexibility into structure-based ligand screening is important in identifying potential HIF-2 binders.

**Figure 1:**
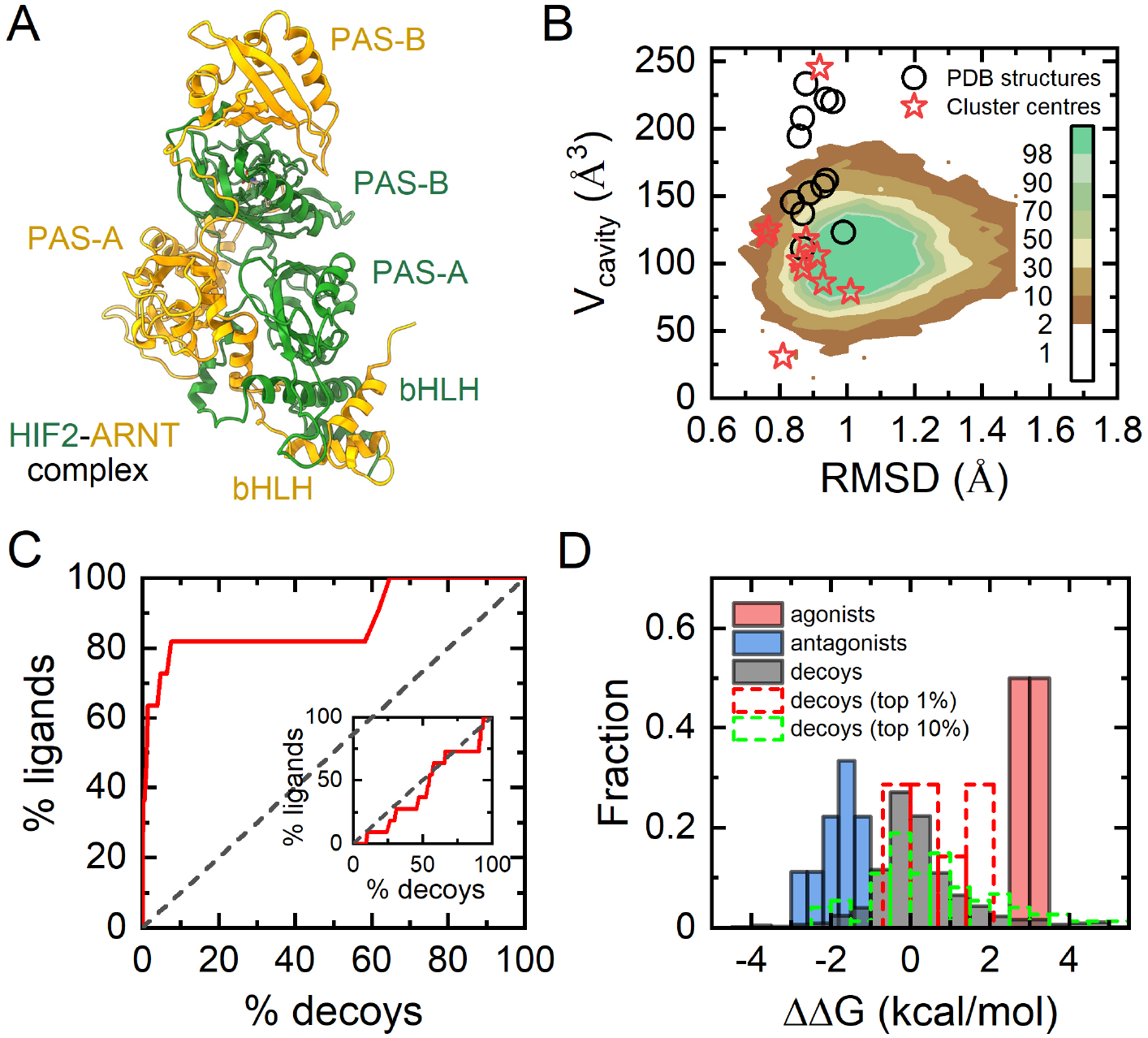
Conformational plasticity of the HIF-2 PAS-B cavity. (A) Ribbon representation of the HIF-2*α*/ARNT complex, with HIF-2*α* shown in green and ARNT in yellow. (B) 2D histogram of the HIF-2 PAS-B domain cavity volume (V_cavity_) and PAS-B C_*α*_ RMSD relative to the experimental apo-state (PDB ID: 4ZP4^35^), computed from 1.5 µs long MD simulations in apo-state. Color represents the sampling frequency. Scattered markers represents V_cavity_ and RMSD corresponding to apo-state cluster centers (stars) and holo-state experimental structures (circles). (C) ROC curve evaluating ligand enrichment over decoys in docking-based virtual screening of the HIF-2 PAS-B domain. Ligands and decoys were ranked according to their docking scores, with the computational binding affinity (Δ*G*) for each ligand defined as the best (lowest) docking score obtained across all PAS-B conformers. Group B conformers (apo-state cluster centers combined with holo-state structures) were used for the main ROC curve, while the inset shows the results using Group A conformers (apo-state cluster centers only) (D) Histogram of ΔΔ*G* (defined in equation 1) values for agonists, antagonists, and decoys.

One practical strategy to incorporate receptor flexibility in docking-based virtual screening is to use an ensemble of diverse binding site conformations.^38^ To assess the adequacy of our selected receptor ensemble and docking parameters, we evaluated their ability to distinguish known ligands from decoys based on rank. ^39^ Two sets of receptor conformations were used: Group A, consisting of 10 cluster centers derived from apo-state MD simulations by clustering frames based on PAS-B cavity shape using POVME 3.0^36^ (details in Supporting Information Sec. 1.4); and Group B, which included all Group A structures along with holo-state conformations from experimentally determined ligand-bound structures of HIF-2 PAS-B (PDB IDs: 3F1O,^29^ 3H82,^30^ 3H7W,^30^ 4GS9,^37^ 4XT2,^40^ 5TBM,^41^ 4ZQD,^35^ 6E3S,^27^ 6E3U,^27^ and 6E3T^27^). We selected a set of 10 ligands (8 antagonists, 2 agonists), listed in Table S1, known to bind the PAS-B cavity and generated 100 property-matched decoys per ligand using the DUDE-Z webserver.^42^ These decoys are compounds with similar physical properties as the known ligands (actives) but have unrelated topologies and are presumed inactive. The decoys for each ligand were generated with following constraints: molecular weight within ±125 g mol^*−*1^, logP within ±3.6, rotatable bonds within ±5, hydrogen bond acceptors (HBA) within ±4, hydrogen bond donors (HBD) within ±3, same charge, maximum Tanimoto similarity between ligand and decoy limited to less than 0.35, and maximum Tanimoto similarity between two decoys to less than 0.8. Docking was performed using Autodock Vina,^43^ and the performance of the multi-conformer set and docking protocol in enriching known ligands over decoys was evaluated using receiver-operating characteristic (ROC) curves, which quantify the true positive rate as a function of the false positive rate. For each ligand, the computational binding affinity (defined as ΔG) was taken as the highest affinity observed among all HIF-2 PAS-B conformers.

Figure 1C shows the ROC curve for the two groups of conformers. Multi-conformer docking with Group A conformers, comprising receptor conformations from the apo-state trajectory only, shows poor enrichment of known ligands over decoys, with an ROC area under the curve (AUC) of 0.47. However, excellent enrichment was obtained with Group B conformers, which additionally include experimental holo-state conformations of the PAS-B domain, yielding a ROC-AUC of 0.87. This performance is comparable to some of the recent successful virtual screening studies.^44–46^ All known ligands and decoys exhibited the highest binding affinity with the holo-state structures, likely due to their larger cavity volume compared to most apo-state cluster centers (Figure S3A). These observations have two distinct explanations. Either, within the conformational selection mechanism, the 1.5 µs MD trajectory of the apo-state is not able to capture full PAS-B plasticity, or the ligand binding to PAS-B domain follows an induced fit mechanism, wherein there is a measurable change in protein conformation on ligand binding. The latter mechanism, either alone or with conformational selection, is expected to be relevant in the present case since the PAS-B domain cavity is buried in the dominant apo-state. Overall, the inclusion of holo-state structures is expected to significantly improve virtual screening accuracy to identify new ligands with high affinity for the HIF-2 PAS-B domain.

For each ligand (except M1002), the best binding affinity was obtained for the docked structure with the corresponding holo-state conformer (see Table S2). For M1002, a known agonist without an available experimental structure, the highest binding affinity was obtained on docking with the HIF-2 PAS-B conformer taken from the M1001-bound state, another known agonist.^27^ Following this observation, we evaluated the difference in binding affinity of a ligand with known agonist-bound and antagonist-bound conformers for identifying a ligand as a decoy, an agonist, or an antagonist: ΔΔG (defined in equation 1).

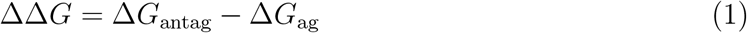

where, Δ*G*_antag_ is the best (lowest) binding affinity of the ligand across all antagonist-bound receptor conformations, and Δ*G*_ag_ is the best (lowest) binding affinity of the ligand across all agonist-bound receptor conformations. This metric gave a very high accuracy in distinguishing known agonists and antagonists for multiple proteins. ^47^ Figure 1D shows that ΔΔG completely separates known HIF-2 agonists from known antagonists, with ΔΔG values for agonists ranging from 2.9 to 3.5 kcal*/*mol, and for antagonists ranging from *−*2.8 to *−*1.3 kcal*/*mol. In contrast, the distribution of ΔΔG for decoys was centered close to 0 kcal*/*mol, but a part of the distribution overlapped with that for the antagonists and ag-onists. We found that 95% and 99% decoys were within ΔΔG of *−*2 to 3.4 kcal*/*mol and *−*3.7 to 4.8 kcal*/*mol, respectively. This overlap significantly decreased when the decoy set was restricted to top 1% on basis of the best obtained binding affinity, with 99% of top 1% decoys within ΔΔG of *−*0.6 to 1.9 kcal*/*mol (complete separation from both agonists and antagonists). Therefore, using ΔΔG as a metric for the enrichment of known ligands over decoys showed excellent performance, with a ROC-AUC of 0.96 for both agonists and antagonists (Figure S3(B-C)). Thus, ΔΔG appears to be an excellent metric for distinguishing agonists from antagonists and both from decoys in virtual screening.

Wu et al. used crystallographic, biophysical, and cell-based functional studies to show that agonist and antagonist binding induced distinct local structural changes at the binding site.^27^ Antagonist binding displaced HIF-2 residue M252 from inside the PAS-B pocket towards the ARNT subunit, weakening heterodimerization. Conversely, agonist binding displaced residue Y281 from inside the PAS-B pocket towards the ARNT subunit, where it formed a hydrogen bond with ARNT residue Y456, thereby strengthening heterodimerization. We defined these intrinsic conformational rearrangements of M252 and Y281 by order parameters *θ*_M252_ and *ϕ*_Y281_, respectively. *θ*_M252_ is the angle made by the vector from C*α* to C*ϵ* of M252 and the vector from C*α* to the PAS-B pocket center (Figure 2A). *ϕ*_Y281_ is the dihedral angle for N, C*α*, C*β*, and C*γ* atoms of Y281 (Figure 2B).

**Figure 2:**
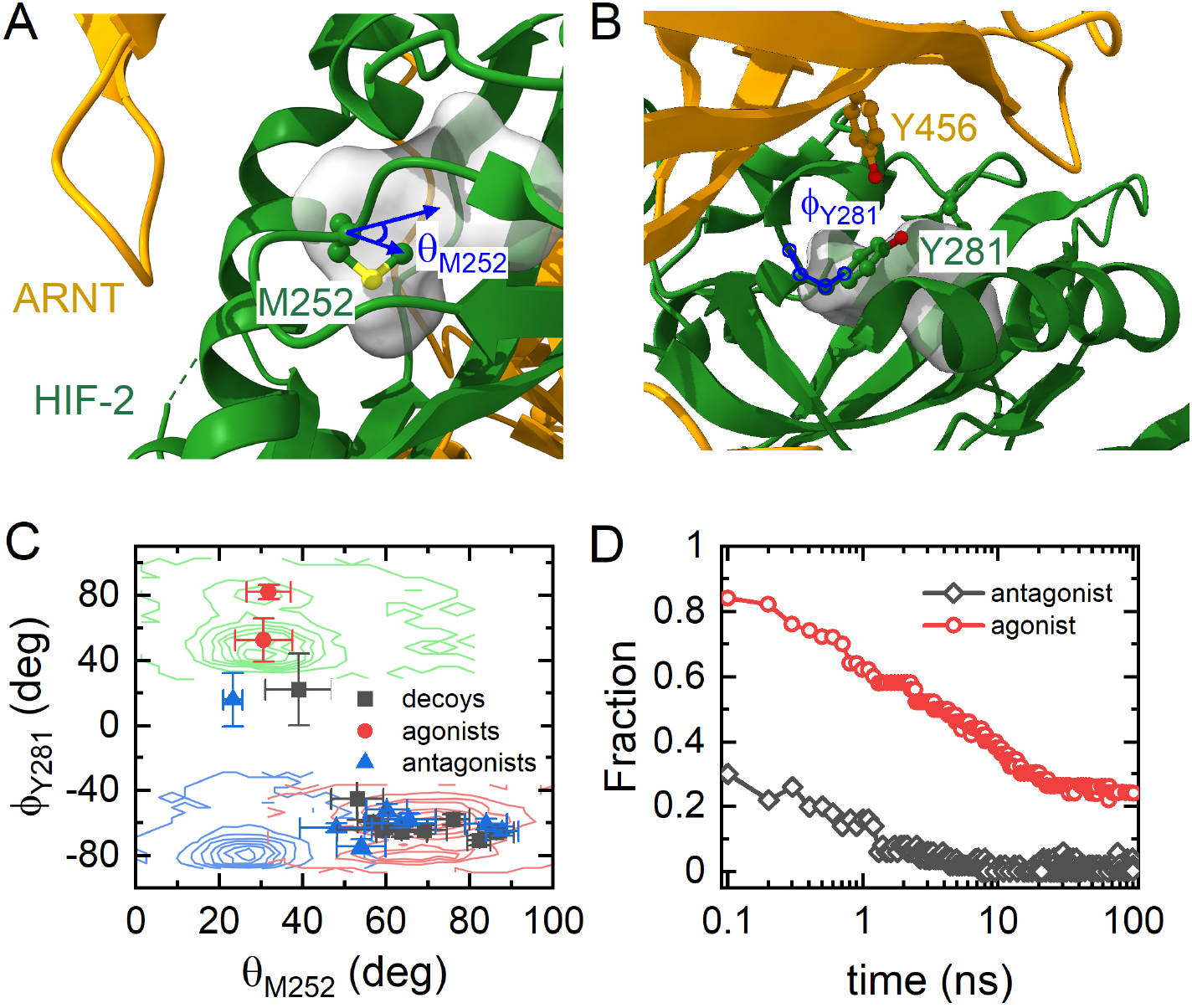
Differential local conformational changes in HIF-2 PAS-B domain upon agonist and antagonist binding. (A) Definition of *θ*_M252_, an angle capturing the displacement of HIF-2 residue M252 from the PAS-B pocket toward the ARNT subunit upon antagonist binding, which weakens heterodimerization. *θ*_M252_ is defined by the angle between two lines: one connecting the C*α* and C*ϵ* atoms of M252, and the other connecting the C*α* atom of M252 to the pocket center. The pocket center is the geometric center of the C*α* atoms of HIF-2 residues 246, 252, 277, 278, 280, 281, 289, 292, 293, 296, 302, 303, 304, 309, 321, and 322. (B) Definition of *ϕ*_Y281_, a dihedral angle that captures the displacement of HIF-2 residue Y281 toward the ARNT subunit upon agonist binding, where it forms a hydrogen bond with ARNT residue Y456, promoting heterodimerization. *ϕ*_Y281_ is defined by the N, C*α*, C*β*, and C*γ* atoms of Y281. (C) Joint distribution of *ϕ*_Y281_ and *θ*_M252_ angles from long (1.5 µs) MD simulations of the apo state (blue contours), antagonist PT2385-bound state (red contours), and agonist M1001-bound state (green contours). Scatter points represent distributions from short (1 ns) MD simulations initiated from the highest-ranked structure obtained via multi-conformer docking using group B conformers for multiple known ligands and highest-ranked decoys. (D) Fraction of MD trajectories out of 50, started from the same initial configuration, in which residue M252 is displaced outwards upon antagonist PT2385 binding (red, circles) and in which residue Y281 is displaced outwards upon agonist M1001 binding (black, diamonds). Simulations were startred from the best configurations obtained from multi-conformer docking using Group A (apo-state conformers only) conformers.

We used long MD simulations (three replicas of 500 ns for each case) of the apo state, holo states with ligands PT2385 (antagonist) and M1001 (agonist) to analyze differences in these order parameters. Holo-state simulations were initiated from the highest-ranked structures obtained from multi-conformer docking of ligand using only Group A (apo-state conformers only) conformers. The ligand topologies were initially generated using CGenFF,^48^ but high penalty scores indicated low confidence in some partial charges. MD simulations using these charges led to significant deviations (~ 2 Å) of the ligand-bound pose from the experimental structure, particularly for PT2385 (Figure S1). To improve charge accuracy, we determined ligand partial charges using density functional theory (DFT) in implicit solvent and QM/MM (quantum mechanics/molecular mechanics) ^49^ calculations with the ligand treated quantum mechanically in the binding pocket. QM/MM-derived charges yielded the lowest deviation (~ 1 Å) from experimental poses for both M1001 and PT2385 and were used for all subsequent simulations (Figure S1). Figure 2C shows the outward displacement of M252 on PT2385 binding, with *θ*_M252_ ~ 30^*°*^ for apo and M1001-bound states while *θ*_M252_ ~ 70^*°*^ for the PT2385-bound state. In contrast, we observed an outward displacement of Y281 on M1001 binding, with *ϕ*_Y281_ *~ −*80^*°*^ for apo and PT2385-bound states while *ϕ*_Y281_ ~ +50^*°*^ for the M1001-bound state. Furthermore, the outward displacement of Y281 upon M1001 binding resulted in weak hydrogen bond formation with ARNT residue Y456: hydrogen-acceptor distance (*d*_HA_) of 3.9±0.7 Å and donor-hydrogen-acceptor angle (*θ*_DHA_) of 91±12^*°*^ (Figure S4). These characteristic local structural changes obtained from MD simulations are in agreement with corresponding crystallographic data on agonist and antagonist binding.^27^

We have noted above that each ligand picked the correct holo state conformer in multiconformer docking experiments. Figure 2C shows that these best holo state structures correctly separate agonists and antagonists in the (*ϕ*_Y281_, *θ*_M252_)-plane. Utility of multiconformer docking was further assessed by carrying short MD simulations of best docked structures obtained when using only Group A (apo-state conformers only) conformers. Figure 2D shows data on the fraction of trajectories (out of 50 independent trajectories) started from ligand-docked apo-state structures, where expected conformational changes, i.e., M252 displacement on PT2385 binding and Y281 displacement on M1001 binding, were observed. For PT2385 binding, correct M252 displacement was observed in 70% of trajectories within 100 ps and in almost 100% of trajectories within 10 ns, implying an induced-fit mechanism for PT2385 (antagonist) binding. On the other hand, correct Y281 displacement was observed in 18% of trajectories within 100 ps while in 23% of trajectories, no displacement is observed in 100 ns. These observations further underscore the importance of multi-conformer docking and inclusion of available holo state structures to identify differential activators of HIF-2 with high computational efficiency.

The analysis of simulation trajectories also helped elucidate the role of ligand binding at HIF-2 PAS B domain on HIF-2/ARNT complex conformational dynamics and DNA binding to this complex. Figure 3A shows that PT2385 binding lead to ~ 50% increase in the per residue root-mean-squared fluctuations (RMSF) of the HIF-2/ARNT complex. In contrast, M1001 slightly decreased the per residue RMSF. Figures S5(A,B) show that there is an increase in RMSF of most of the residues upon PT2385 binding, including the residues far from the ligand-binding site, such as the DNA-binding bHLH residues. Conversely, M1001 binding results in unchanged fluctuations for most residues, while a few regions show decreased fluctuations. This suggest a strong allosteric network in the complex that can be modulated by ligand binding at the HIF-2 PAS-B domain. These findings correlate well with the experimentally observed strong antagonistic properties of PT2385 and the weak agonism of M1001. Notably, these effects on flexibility become apparent only after a minimum simulation timescale of 200-300 ns, highlighting the need for sufficiently long simulation trajectories to capture these responses. The effect of ligand binding on heterodimer flexibility is also expected to affect stability, here quantified using MM/PBSA (Molecular Mechanics Poisson Boltzmann Surface Area) binding energy^50^ (method details in Supporting Information Sec. 1.5). The MM/PBSA absolute binding free energy typically represent an overestimate because of the use of implicit solvent model and neglect of protein conformational entropy, wherein the latter is expected to be particularly important for large protein–protein complexes. Nevertheless, it remains a valuable metric for assessing relative changes in binding affinities across comparable systems.^51,52^ Figure 3B shows that PT2385 binding destabilized the HIF-2/ARNT by 306 kJ mol^*−*1^ (w.r.t. apo state), whereas M1001 binding had a negligible effect on binding energy.

**Figure 3:**
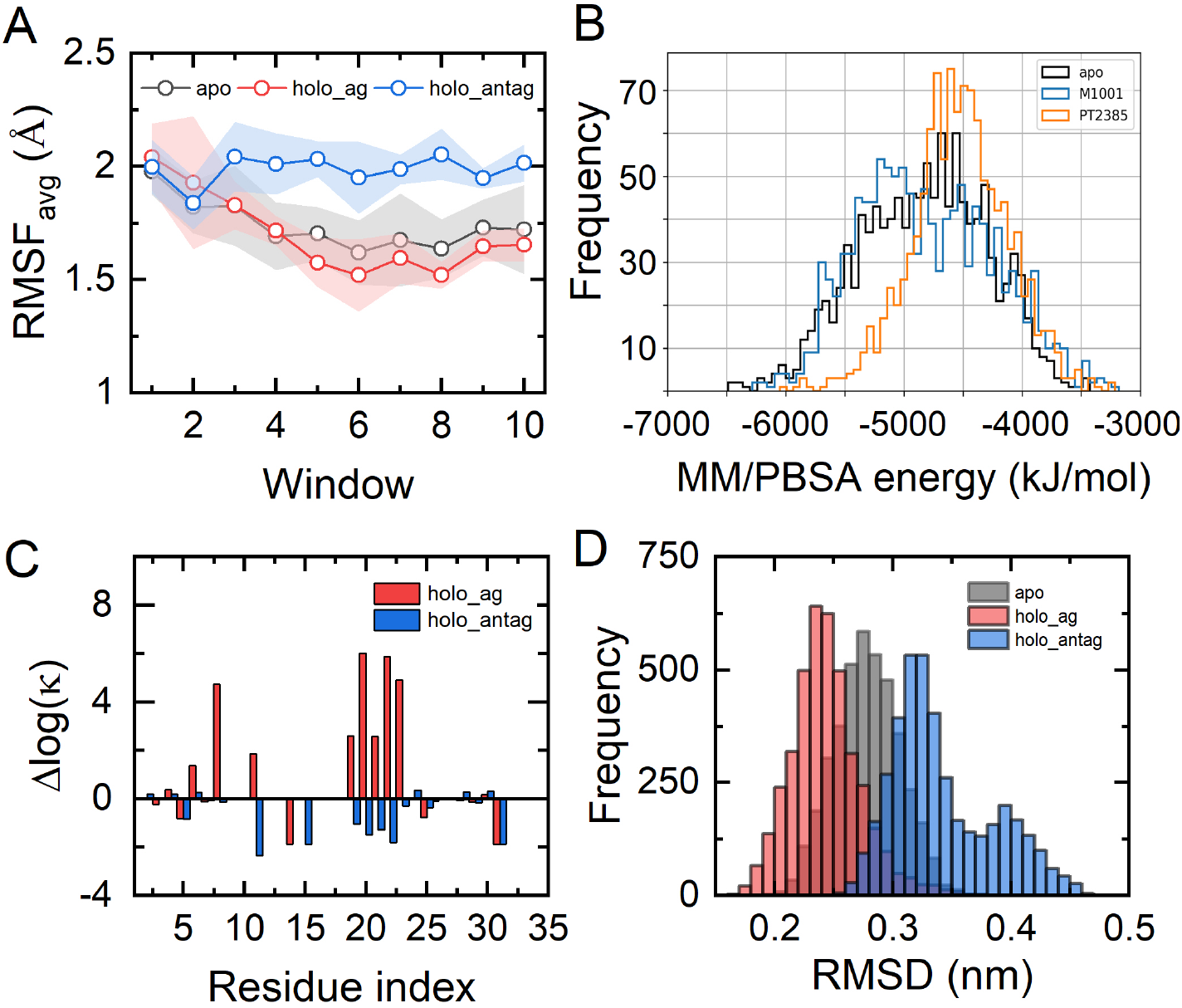
Effect of ligand binding on global structural changes, flexibility, and stability of the HIF2-ARNT complex. (A) RMSF_avg_ (average RMSF of all residues in the HIF-2–ARNT complex) computed from a series of 50 ns windows within a total of 500 ns long simulations. Shaded region represents standard error in the estimates of RMSF_avg_. (B) Histogram of MM/PBSA binding energy between HIF-2 and ARNT for apo, PT2385-bound, and M1001-bound states.(C) Change in the amide hydrogen protection factors, *κ* (defined in equation S2), in the holo state relative to the corresponding apo-state values for residues in the DNA-binding domain (DBD), estimated from MD simulations. Residue indices 1-17 correspond to HIF-2 bHLH domain residues 28-44, and indices 18-35 correspond to ARNT bHLH domain residues 100-117 (D) Histogram of DBD C_*α*_ RMSD in a given state relative to the DNA-bound conformation of the DBD.

The changes in the HIF-2/ARNT interface stability is currently considered the primary mechanism by which these ligands exert their effect. ^27,29,31^ Here, we determined the effect of ligand binding on the structure and dynamics of the DNA-binding bHLH domains and its connection to transcriptional activity. Specifically, we calculated amide protection factors, *κ*, defined as the ratio of the probability of a residue being in a closed state, where hydrogendeuterium exchange (HDX) is not possible, to the probability of being in an open state (details in the Supporting Information Sec. 1.6). Figure 2C shows that M1001 binding increased the protection of bHLH residues, particularly those in the ARNT bHLH domain, while PT2385 binding reduced the protection of these residues. This implies an entropic effect of ligand binding at the PAS-B domain at the distant DNA-binding domain (DBD), wherein increased flexibility of DBD on antagonist binding is expected to weaken the DNA binding. Figure 2D shows the C_*α*_ RMSD of the DNA-binding domain (DBD) (in apo and holo states) relative to the DNA-bound conformation of the heterodimer (shown in Figure S6A). M1001 (agonist) binding resulted in a DBD conformation closer to that required for DNA binding, whereas PT2385 (antagonist) binding produced the opposite effect. For decoys, the RMSD remained similar to the apo-state. Therefore, in addition to the entropic effect, ligand binding at the PAS B domain also exerts a conformational change at the DBD, making it either more similar (in case of agonist) or less similar (in case of antagonist) to the DNA-bound conformation when compared with the DBD conformation in the apo state of HIF-2/ARNT complex. These findings clearly show that antagonist binding at the HIF-2 PAS B domain allosterically alters the structure and dynamics of the DBD, weakening its interactions with the hypoxia response element (HRE). Our findings are in excellent agreement with the observed effect of PT2385 and M1001 binding on residue flexibility determined using HDX mass spectrometry and the weakening of heterodimer binding with the HRE as determined using cell-based activity assays.^27^ We recommend that determination of the overall effect of ligand binding on HIF-2/ARNT complex structure and dynamics, both at the heterodimer binding interface and the DNA-binding bHLH domain, will be crucial in distinguishing agonists from antagonists and actives from decoys.

Further distinctions between agonists and antagonists are observed in terms of local conformational changes. Antagonist binding displaces the HIF-2 residue M252 from inside the PAS-B pocket towards the ARNT subunit, weakening heterodimerization. In contrast, agonist binding displaces residue Y281 from the same pocket towards the ARNT subunit, where it forms a hydrogen bond with ARNT residue Y456, thereby strengthening heterodimerization. This dynamic requires extensive MD simulations to analyze and serves as a crucial descriptor to distinguish between agonists and antagonists. Additionally, agonists and antagonists have differing effects on the overall conformational fluctuations of the protein. Antagonists increase these fluctuations, potentially leading to destabilization, whereas agonists decrease them. The HIF-2/ARNT interface binding energy also serves as an important descriptor, with antagonists generally decreasing interface stability and agonists increasing it. Finally, agonists and antagonists affect the DNA-binding bHLH domain’s structure and dynamics differently. Antagonist binding displaces the bHLH domain structure away from the configuration required for binding with the hypoxia response element (HRE), while agonist binding enhances alignment with the HRE. Additionally, antagonist binding increases the flexibility of bHLH residues, potentially resulting in a higher entropic penalty upon DNA binding and thus reduced affinity. These descriptors collectively provide a robust framework for prioritizing molecules in virtual screening and should be considered when selecting ligands for experimental evaluation.

In summary, our study demonstrates that known ligands and decoys distinctly modulate the structure and dynamics of the HIF-2/ARNT complex. These differences can be leveraged to computationally distinguish agonists from antagonists and effectively identify potential actives from decoys in virtual screening, thereby reducing false positives. The HIF-2 PAS-B domain, which serves as the ligand-binding site, exhibits significant conformational plasticity. Accounting for this flexibility is critical for accurate docking-based virtual screening, and we show that a multi-conformer docking approach, particularly using holo-state experimental structures, improves ligand enrichment over decoys. Further, the differential ΔΔG values effectively differentiate known HIF-2 agonists, peaking around ΔΔG = +3 kcal*/*mol, from antagonists, which peak at ΔΔG = *−*2 kcal*/*mol. This metric is computationally efficient as it does not require MD simulations and can be computed for a large number of ligands. These metrics can be used as initial screening strategy to enable prioritization of a manageable subset of candidates, on the order of hundreds, for subsequent analysis using more computationally intensive MD-based descriptors. These include ligand-induced local conformational changes, such as the displacement of M252 (by antagonists) or Y281 (by agonists) within the PAS-B pocket. Antagonist-induced displacement of M252 weakens HIF-2/ARNT heterodimerization, while agonist-induced displacement of Y281 promotes heterodimer stabilization via a hydrogen bond with ARNT Y456. Further, agonists and antagonists differentially modulate global conformational fluctuations: antagonists generally increase flexibility, potentially destabilizing the complex, whereas agonists reduce fluctuations, favoring stability. The interface binding energy between HIF-2 and ARNT also serves as a key descriptor, with agonists enhancing and antagonists reducing interface stability. Importantly, these ligands differently influence the structure and dynamics of the DNA-binding bHLH domain. Antagonists disrupt alignment with the hypoxia response element (HRE) and increase bHLH flexibility, whereas agonists enhance alignment and reduce flexibility, likely improving DNA-binding affinity. Together, these descriptors form a robust, multiscale framework for ligand prioritization in HIF-2-targeted virtual screening. We are currently applying this strategy in combination with AI-aided virtual screening to explore a chemical library of approximately 500 million molecules for the discovery of novel agonists and antagonists for selective modulation of HIF-2 transcriptional activity.

## Notes

### Competing Interest Statement

The authors have declared no competing interest.

