## Supporting Information for "Structural, Thermodynamic, and Dynamic Descriptors for Differential Mechanism of HIF-2 Activity Modulators"

### 1 Supplementary Methods

#### 1.1 Docking Calculations

Molecular docking was performed using AutoDock Vina.[1] A grid box of size 25 Å along each of the $x$, $y$, and $z$ dimensions was used, with the grid centered at the geometric center of the C$\alpha$ atoms of the following HIF-2 PAS-B domain residues: 246, 252, 277, 278, 280, 281, 289, 292, 293, 296, 302, 303, 304, 309, 321, and 322. The exhaustiveness parameter was set to 8 for all docking runs. For each ligand, docking was performed against an ensemble of HIF-2 PAS-B conformers. Two receptor ensembles were used: (i) **Group A**, consisting of cluster centers from apo-state molecular dynamics simulations, and (ii) **Group B**, comprising Group A conformers along with holo-state structures from available experimental complexes. The binding affinity for each ligand was defined as the best (lowest) docking score obtained across all conformers within the respective ensemble.

#### 1.2 All-Atom Molecular Dynamics Simulations

MD simulations of the HIF-2/ARNT complex were conducted using GROMACS 2021.4[2] with the CHARMM36 force field.[3] The initial configuration was taken from the PDB database[4] (PDB ID 4ZP4[5]). Missing residues were modelled using AlphaFold2. The initial structure (residues 100-464 of ARNT and 28-360 of HIF-2) was centred in a cubic box with a minimum distance of 1 nm between the protein and the box edge. The system was solvated with the TIP3P water molecules and the charge was neutralized with $Na^+$ and $Cl^-$ ions, adjusting the ionic strength to 0.1 M. Energy minimization was performed using the steepest descent algorithm with a maximum force criterion of less than 100 kJ/mol to remove potential steric clashes. Following minimization, the system was equilibrated in the NVT ensemble at 310 K using the velocity rescale thermostat, ensuring temperature stability. This was followed by pressure equilibration in the NPT ensemble at 310 K and 1 bar using the C-rescale barostat to stabilize the system's density. For the MD simulations, three 500 ns replicas were executed for both apo and holo states of the complex to ensure adequate sampling. Long-range electrostatic interactions were treated using the Particle Mesh Ewald method, and a cutoff of 1.2 nm was used for both Coulombic and van der Waals interactions. Temperature and pressure were controlled using the modified velocity-rescale thermostat and C-rescale barostat, respectively. Root mean squared fluctuations (RMSF) of C$\alpha$ atoms of residues was calculated using the GROMACS command *gmx rmsf*, after alignment of each simulation trajectory frame to the average structure of the simulation window to remove overall translation and rotation.

##### 1.3 Ligand Force Field Parameterization

For the MD simulations of PT2385-bound and M1001-bound states, initial ligand parameters were determined using the CGenFF[6] via CHARMM-GUI.[7] However, some of the estimated partial charges on the atoms had high penalties or low confidence scores. Utilizing these inaccurate charges could result in significant deviations in the ligand binding mode from the experimentally observed mode during MD simulations (Figure S1). To rectify this issue, we reparameterized the ligand atom charges using two approaches: Density Functional Theory (DFT) and Quantum Mechanics/Molecular Mechanics (QM/MM).

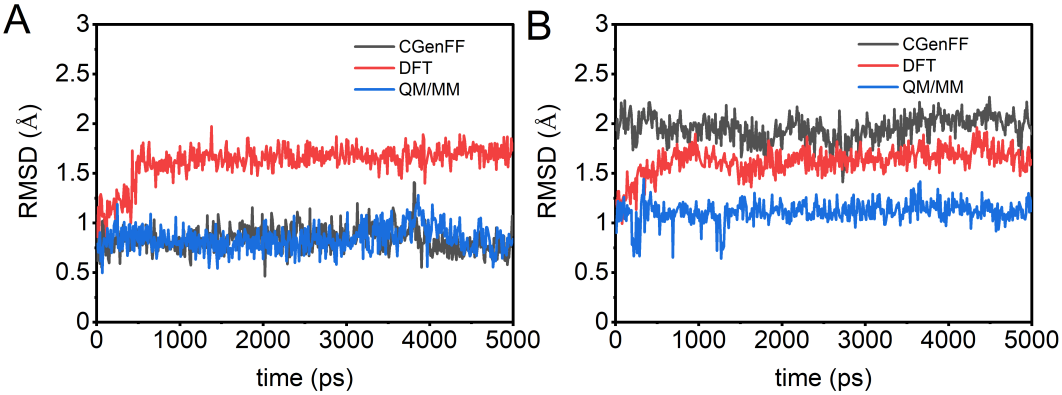


Figure S1: Ligand RMSD relative to the initial state in MD simulations performed with charges determined using CGenFF, DFT, and QM/MM for (A) M1001 and (B) PT2385.

##### 1.3.1 DFT Charges:

The DFT charges were calculated by first optimizing the geometry of the ligand in an implicit solvent using the COSMO model. The B3LYP exchange-correlation functional and the 6-31G** basis set were employed for this optimization. Partial charges were then obtained through ESP-fitting of the electron density. Despite these efforts, simulations using these DFT-derived charges still showed deviations in the ligand binding mode (Figure S1).

##### 1.3.2 QM/MM Charges:

We subsequently adopted a QM/MM approach using GROMACS coupled with CP2K,[8] where the ligand was treated at the quantum mechanics (QM) level, while the protein and explicit water molecules were treated at the molecular mechanics (MM) level. The QM/MM parameters included the GPW (Gaussian and Plane Waves) method, a TZV2P (triple-zeta valence with 2 polarization functions) basis set, an energy cutoff of 450 eV, GGA-PBE exchange-correlation functional, DFT-D3 dispersion correction, EPS_SCF $= 5 \times 10^{-8}$, CHARMM36 force field, and RESP fitting for charge calculation. This approach resulted in the preservation of the ligand binding mode, with RMSD values $\lesssim 1$ Å during MD simulations (Figure S1). The QM/MM-derived charges provided a more accurate representation of the ligand's behavior in the binding pocket.

#### 1.4 Shape-based clustering of PAS-B cavity

Shape-based clustering of the HIF-2 PAS-B binding pocket conformations was performed using POVME 3.0.[9] The MD simulation trajectory was processed with POVME to calculate a pairwise pocket similarity matrix across all frames, using the Tanimoto overlap score as the similarity metric. The Tanimoto score was computed based on the overlap of pocket grid points. The score between two pocket conformations was calculated as the number of shared grid points divided by the total number of grid points present in either conformation. The resulting similarity matrix was used for k-means clustering of pocket shapes, with the number of clusters set to 10.

##### 1.5 MM/PBSA binding energy of HIF2-ARNT complex

The MM/PBSA binding energy was computed for every 10ps frame of a MD simulation trajectory using *g_ mmpbsa*.[10] The MM/PBSA parameters used are provide below (for details about the keywords and units, refer to the *g_ mmpbsa* documentation).

```
polar   = yes                      swin   = 0.30
cfac    = 3.0                      sdens   = 10
gridspace  = 0.5                  temp    = 330
fadd    = 30                       bcfl   = mdh
gmemceil  = 32000                 PBsolver  = npbe
pcharge  = 1                      apolar  = yes
prad  = 0.95                       gamma        = 0.0226778
pconc        = 0.100              sasrad       = 1.4
ncharge  = -1                     sasaconst    = 3.84982
nrad  = 1.81                       bconc  = 0.033428
nconc   = 0.100                    dpos      = 0.05
pdie    = 2                        APsdens  = 20
sdie    = 80                       grid         = 0.45 0.45 0.45
vdie    = 1                        APsrfm       = sacc
srad    = 0.6                      APswin       = 0.3
chgm         = spl4               APtemp       = 300
srfm         = smol
```

##### 1.6 Amide Hydrogen Protection Factors

Hydrogen/Deuterium exchange (H/D-ex), monitored by NMR or mass spectrometry (MS) is widely employed to protein flexibility and dynamics.[11–13] H/D-ex can only occur if the amide is exposed to solvent, therefore suggesting resdiue fluctuations an integral part of the exchange mechanism. According to the standard H/D-ex model,[14] an amide can exist in a closed (C) state where no exchange occurs, or in an open (O) state where exchange occurs at a rate $k_{\text{int}}$ (see equation S1).

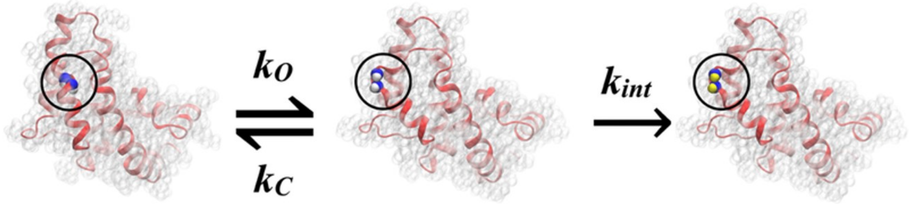


Figure S2: Kinetic model for hydrogen/deuterium exchange. Conformational shifts expose internal amide groups (blue) (closed state) to the solvent (open state), allowing the exchange of amide hydrogens (white) with deuterium (yellow) at an intrinsic rate constant $k_{int}$. Figure adapted from Ref.[15]

$$(\text{N}-\text{H})_\text{C} \underset{\text{k}_\text{cl}}{\overset{\text{k}_\text{op}}{\rightleftharpoons}} (\text{N}-\text{H})_\text{O} \xrightarrow{\text{k}_\text{int}} (\text{N}-\text{D})_\text{O} \tag{S1}$$

Assuming, the rate of H/D-ex from the O state is same as in model peptides with the same neighboring side chains, i.e $k_{\text{int}} = k^0_{\text{HX}}$, the observed differences in overall H/D-ex can be attributed to the change in amide protection factors, $\kappa$, where $\kappa = k_{\text{cl}}/k_{\text{op}}$. Several models[15–18] exists that empirically relates $\kappa$ to properties of amide such as buried surface area,[15] number of hydrogen bonds,[16–18] number of residues in the vicinity[16] etc., that can be obtained from MD simulations. Alternatively, at equilibrium, $\kappa$ relates to the fractional population of O and C states, $f_O$ and $f_C$ respectively, as $\kappa = f_C/f_O$ from detailed balance. Persson and Halle[19] proposed that amides exchange hydrogen with water molecules in locally distorted transient conformation with two water molecules directly coordinated to the N–H group. Thus, the O state was defined as the one where amide hydrogen has atleast 2 water oxygens with 2.6 Å radius (precise radius value is not critical, see detailed discussion in Ref.[19]).

$$\kappa = \frac{f_C}{f_O} = \frac{N_C}{N_O} \tag{S2}$$

where, $N_O$ and $N_C$ is the number of trajectory frames where amide is in open and closed state respectively. This method has be shown to reproduce experimental protection factors with better accuracy than existing empirical methods for multiple proteins.[15] Statistical uncertainties in calculations of $\kappa$, due to finite length of MD simulation trajectories, is estimated by equation S3.[19]

$$\sigma(\kappa) = \kappa\sqrt{\frac{1}{N_{\mathrm{C}}} + \frac{1}{N_{\mathrm{O}}}} \tag{S3}$$

#### 2 Supplementary Figures

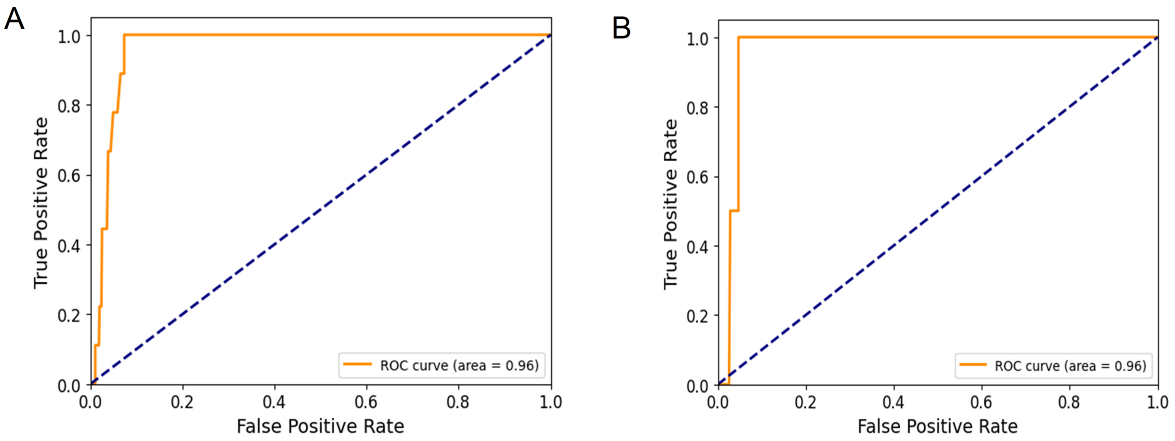


Figure S3: ROC curves obtained from ranking ligands and decoys based on increasing $\Delta\Delta G$ for antagonists (A) and decreasing $\Delta\Delta G$ for agonists (B).

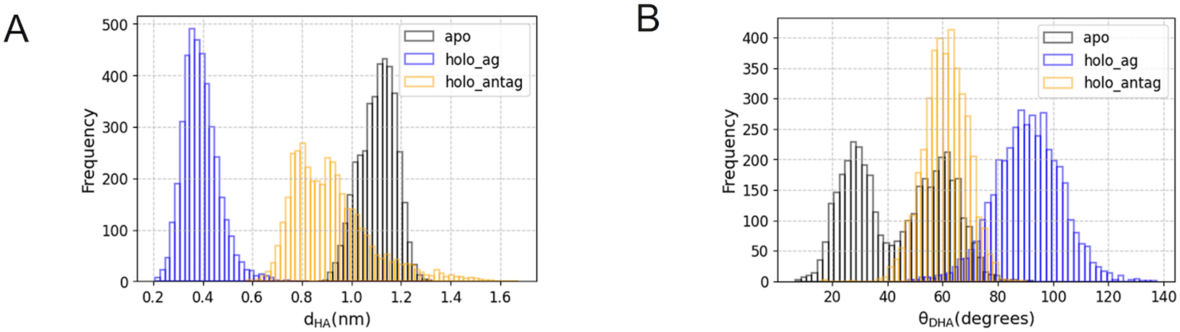


Figure S4: Distribution of hydrogen-acceptor distance $d_{\mathrm{HA}}$ (C) and donor-hydrogen-acceptor angle $\theta_{\mathrm{DHA}}$ (D) to assess hydrogen bond formation between HIF-2 residue Y281 and ARNT residue Y456.

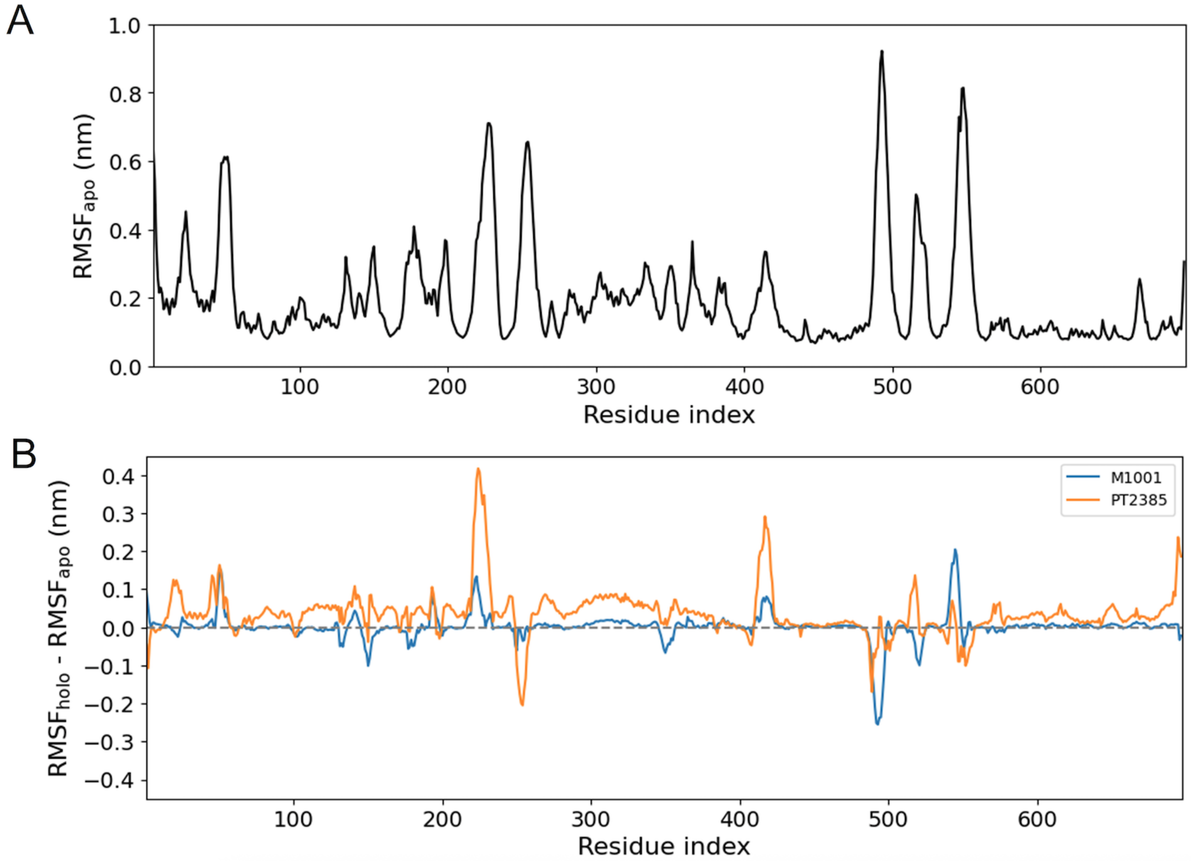


Figure S5: (A) Residue $C_\alpha$ root mean squared fluctuations (RMSF) profile of the HIF-2/ARNT complex in the apo-state. The mean RMSF value of each residue is calculated from three independent 500 ns MD trajectories. Residue indices 1-365 correspond to ARNT residues 100-464, and residue indices 366-698 correspond to HIF-2 residues 28-360. (B) Changes in the average RMSF profile upon ligand binding.

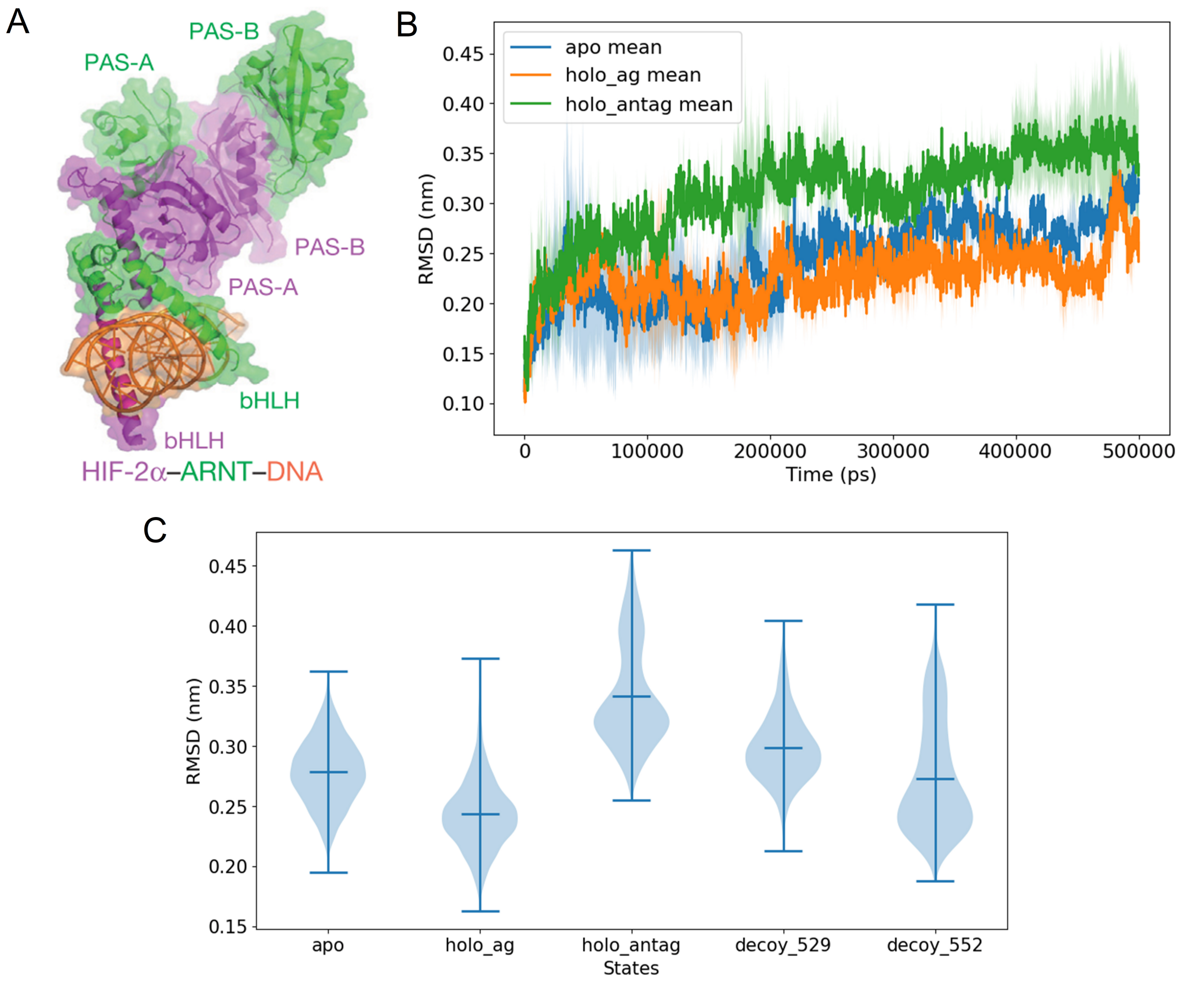


Figure S6: (A) Crystal structure of the HIF-2/ARNT complex with hypoxia response element (HRE) DNA, obtained from PDB accession code 4ZPK.[5] (B) Mean $C_\alpha$ RMSD of DNA-binding bHLH domains relative to the 4ZPK structure, computed from three independent replicas of 500 ns MD trajectories for apo, PT2385-bound (holo_antag), and M1001-bound (holo_ag) states. The shaded region represents standard deviations. (C) Distribution of $C_\alpha$ RMSD of DNA-binding bHLH domains relative to the 4ZPK structure for apo, active ligand-bound, and decoy-bound states.

#### 3 Supplementary Tables

**Table S1: Ligands known to experimentally bind the HIF-2$\alpha$ PAS-B domain. M1001 and M1002 are weak agonists, while the remaining ligands are antagonists.**

| Ligand | Structure | Ref. | Ligand | Structure | Ref. |
|---|---|---|---|---|---|
| THS-044 | 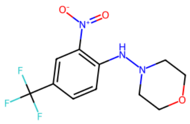 | 20 | THS-020 | 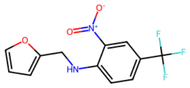 | 21 |
| THS-018 | 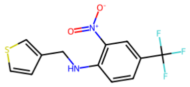 | 21 | 0XB | 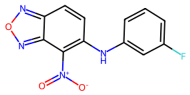 | 22 |
| 0X3 | 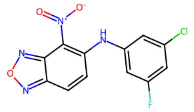 | 5 | 43L | 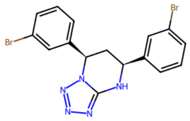 | 23 |
| T1001 | 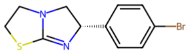 | 13 | PT2385 | 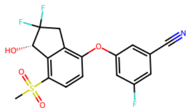 | 13 |
| M1001 | 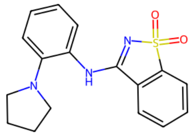 | 13 | M1002 | 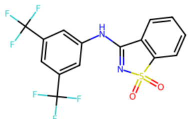 | 13 |

**Table S2: Docking binding affinities $\Delta G$ (in kcal/mol) of ligands against multiple receptor conformers.**

| **Ligand** | **3F1O**[20] | **3H82**[21] | **3H7W**[21] | **4GS9**[22] | **4XT2**[23] | **5TBM**[24] | **4ZQD**[5] | **6E3S**[13] | **6E3U**[13] | **6E3T**[13] |
|---|---|---|---|---|---|---|---|---|---|---|
| THS-044 | **-9.0** | -8.4 | -7.9 | -8.8 | -8.0 | -8.5 | -8.5 | -7.9 | -6.9 | -7.5 |
| THS-020 | -7.7 | **-8.2** | -7.9 | -7.9 | -7.9 | -8.0 | -7.6 | -6.8 | -6.6 | -7.4 |
| THS-018 | -8.0 | -8.1 | **-8.1** | -7.4 | -7.7 | -6.9 | -7.2 | -7.2 | -6.0 | -7.1 |
| 0XB | -6.8 | -7.2 | -7.7 | **-7.9** | -7.7 | -7.4 | -7.6 | -7.5 | -6.6 | -6.9 |
| 0X3 | -7.5 | -8.2 | -8.3 | -7.9 | -8.0 | -8.6 | **-8.8** | -8.0 | -6.0 | -7.4 |
| 43L | -7.0 | -7.7 | -7.9 | -7.2 | **-8.0** | -7.3 | -7.2 | -7.5 | -5.7 | -7.5 |
| T1001 | -7.4 | -7.1 | -8.0 | -8.0 | -7.2 | -7.4 | -7.2 | -7.8 | -6.4 | **-8.2** |
| PT2385 | -7.8 | -7.6 | -7.1 | -6.9 | -8.0 | -8.4 | -7.7 | **-8.5** | -6.7 | -7.9 |
| M1001 | -5.0 | -4.9 | -5.0 | -5.1 | -6.6 | -6.4 | -6.2 | -6.1 | **-10.1** | -6.5 |
| M1002 | -5.1 | -5.1 | -5.8 | -5.5 | -6.4 | -7.0 | -6.3 | -5.9 | **-9.9** | -6.3 |

(6) Vanommeslaeghe, K.; Hatcher, E.; Acharya, C.; Kundu, S.; Zhong, S.; Shim, J.; Darian, E.; Guvench, O.; Lopes, P.; Vorobyov, I. et al. CHARMM General Force Field

(CGenFF): A force field for drug-like molecules compatible with the CHARMM all-atom additive biological force fields. *Journal of computational chemistry* **2010**, *31*, 671.

(13) Wu, D.; Su, X.; Lu, J.; Li, S.; Hood, B. L.; Vasile, S.; Potluri, N.; Diao, X.; Kim, Y.;

Khorasanizadeh, S. et al. Bidirectional modulation of HIF-2 activity through chemical ligands. *Nature Chemical Biology 2019 15:4* **2019**, *15*, 367–376.
